# Single-cell analysis of cell fate bifurcation in the chordate *Ciona*

**DOI:** 10.1101/2020.10.19.346213

**Authors:** Konner M. Winkley, Wendy M. Reeves, Michael T. Veeman

## Abstract

Inductive signaling interactions between different cell types are a major mechanism for the further diversification of embryonic cell fates. Most blastomeres in the model chordate *Ciona robusta* become restricted to a single predominant fate between the 64-cell and mid-gastrula stages. We used single-cell RNAseq spanning this period to identify 53 distinct cell states, 25 of which are dependent on a MAPK-mediated signal critical to early *Ciona* patterning. Divergent gene expression between newly bifurcated sibling cell types is dominated by upregulation in the induced cell type. These upregulated genes typically include numerous transcription factors and not just one or two key regulators. The Ets family transcription factor Elk1/3/4 is upregulated in almost all the putatively direct inductions, indicating that it may act in an FGF-dependent feedback loop. We examine several bifurcations in detail and find support for a ‘broad-hourglass’ model of cell fate specification in which many genes are induced in parallel to key tissue-specific transcriptional regulators via the same set of transcriptional inputs.

## Introduction

The journey from the totipotent fertilized egg to the myriad distinct cell types found in mature animals involves a series of decision points in which cells can follow different trajectories of differentiation [1]. It is now known that these cell fate bifurcations sometimes involve the asymmetric inheritance of cell fate determinants between sibling cells, but more often involve inductive interactions between different cell types. Decades of work in various model organisms have mapped out inductive interactions between many different cell types, identified specific signal transduction pathways used to induce specific cell fates, and identified important transcription factors controlling cell type-specific gene expression. Until recently, however, the global transcriptomic changes underlying bifurcations in cell fate have been elusive. RNAseq can be used to transcriptionally profile cell populations purified by FACS, but it is hard to scale this strategy to profile the full complexity of cell types in even simple developing embryos. Also, the maturation times for fluorescent markers of cell fates make it hard to purify distinct cell types until well after they are first specified. With the advent of massively parallel single cell RNA sequencing (scRNAseq) [2–6], however, it is now possible to reconstruct cell-type specific transcriptional profiles from heterogenous cell mixtures including entire dissociated embryos. scRNAseq has been used to produce atlases of cell-type specific gene expression at key stages in several model organisms [7–10].

### Early development in *Ciona*

In order to quantify the transcriptomic changes associated with cell fate bifurcation events, it is necessary to identify mother-daughter-sibling relationships between scRNAseq clusters. Here we use the mother-daughter-sibling terminology to refer to relationships between transcriptional states as one cell type bifurcates into two. This will vary in different contexts in how congruent it is with the actual lineages of cell divisions. Identifying these relationships de novo presents a complicated technical and conceptual challenge and has been the primary focus of many scRNAseq studies [11–15]. This is especially true in systems where differentiation is not synchronized and pseudotime inference strategies are needed to reconstruct developmental trajectories [16,17]. In the invertebrate ascidian chordate *Ciona,* however, the early lineages are completely stereotyped (Figure S1), development is highly synchronous, and extensive fate mapping and traditional gene expression studies provide extensive and near-comprehensive prior information about the expected cell types and lineage relationships between them as reviewed in [18]. *Ciona* embryos are small and simple, yet stereotypically chordate. This makes relatively deep coverage of each cell type for an entire chordate embryo easily achievable in single-cell experiments. Most ascidian blastomeres become restricted to a single fate during a narrow window between the 64-cell and mid-gastrula stages [19–21], allowing the analysis of many cell fate specification events with a relatively short time course of sequenced stages. The mid-gastrula stage is reached in only ~6hrs with most blastomeres having gone through 8 cell cycles. Development from the fertilized egg to the hatched tadpole larva takes less than 24 hours. Several other studies have used scRNAseq to address diverse questions in ascidian models [10,22–26], but here we provide the first scRNAseq analysis of the critical 64-cell to mid-gastrula time period in *Ciona robusta.*

The early patterning of the bilaterally symmetrical ascidian embryo up to the 32-cell stage relies primarily on two intersecting patterning systems. The first involves cortical rearrangements of the ooplasm and polarity breaking events in response to sperm entry, which culminate in the formation of a structure known as the centrosome attracting body (CAB) in the posterior vegetal blastomeres at the 8-cell stage [27]. The CAB is then continuously partitioned into the posterior-most vegetal cells during subsequent asymmetric divisions and is thought to be responsible for directly or indirectly driving most of the anterior-posterior patterning of the early embryo (reviewed in [28]). The second system involves reciprocal interactions between maternally deposited Gata.a factors and nuclear β-catenin signaling activated on the vegetal side of the early embryo. In subsequent rounds of division at the 16-cell and 32-cell stages, these two pathways lead to the establishment of the three germs layers through antagonistic gene expression and restricted domains of nuclear localization [29–32].

At the 32-cell stage only a subset of the endodermal blastomeres are restricted to a single fate, but nearly all the remaining blastomeres become fate restricted in the next two cell cycles [19–21]. The majority of the fate bifurcation events that take place in this time window are dependent on MAPK signaling [33–38]. MAPK activity at these stages is controlled by FGF ligands expressed on the vegetal side of the embryo downstream of β-catenin [39,40], and by the FGF antagonist ephrinAd, which is expressed in the animal hemisphere downstream of Gata.a [36]. As development continues through the 110-cell and mid-gastrula phases, other FGF agonists and antagonists become expressed in other lineages such as the trunk lateral cells, and in complex patterns in the neural plate [41].

MAPK signaling directly induces several cell fates but also has well-characterized indirect effects. MAPK signaling induces expression of a Nodal ligand in a lateral animal cell population [42]. The Nodal signal activates downstream expression of Notch pathway ligands in neighboring cells and this Nodal/Delta relay further refines several tissue types at the 110-cell and mid-gastrula stages. This is particularly evident in the mediolateral patterning of the neural plate [42].

MAPK activity and repression are thought to be either directly or indirectly responsible for nearly all the fate bifurcations occurring after the 32-cell stage in the *Ciona* embryo [34,36,37,41,43,44]. We performed whole embryo scRNAseq at three key stages during cell fate specification both with and without the MEK inhibitor U0126. Using this data, we set out to characterize the repertoire of transcriptomic states in the early *Ciona* embryo and to define the transcriptomic responses of different precursor cells in response to FGF/MAPK signaling. We exploited the fixed lineages of *Ciona* to ask broad, systems-level questions about the gene regulatory networks (GRN) that control cell fate specification events, with an emphasis on inferring new features of the GRN controlling notochord specification and differentiation.

## Results and Discussion

### Single-cell RNAseq of *Ciona* embryos recapitulates known expression patterns and reveals new transcriptional states

We performed single-cell RNA sequencing of dissociated *Ciona* embryos using a modified dropSeq approach [3] at the 64-cell, 110-cell, and mid-gastrula stages (stages 8, 10, and 12 in Hotta’s *Ciona* staging series [45]) (Figure 1A). These stages are each spaced less than an hour apart at 18°C and span a period in which most *Ciona* blastomeres become restricted to a single major fate. In order to obtain enough cells for each timepoint to ensure deep coverage of each left-right blastomere pair, we pooled gametes from multiple hermaphroditic adults for each fertilization. The small and well-defined number of bilaterally symmetrical cell pairs at these stages provides an upper limit on the number of distinct cell types that are theoretically possible. We profiled over 6,000 cells in wild-type embryos across the three stages and achieved an average coverage of 42 times per theoretical blastomere pair. Most cell types are made up of more than one blastomere pair, so actual coverage of distinct cell types was typically higher. The scRNAseq libraries were sequenced to an average depth of 23,985 reads per cell, representing an average of 11,706 transcripts detected in each cell (Full sequencing details in Supplemental Table 1). Upon initial post-sequencing analysis and clustering, we generated first-pass UMAP plots. Each point on the UMAP plots represents a single set of 3’ RNAs associated with a unique cell barcode (also known as a “single-cell transcriptome attached to microparticle” (STAMP)), and the distances between points represent similarities and differences in gene expression across the entire transcriptome. The distinct clusters evident on the UMAP plots represent different cell types with their own distinct transcriptional states. We compared the gene expression profiles of the STAMPs contained within each cluster to those already known from extensive in situ hybridization screens in *Ciona* [46–48]. We identified several duplicate clusters for many cell types (Figure S2 A-C). These were unexpected based on previous *Ciona* literature, and upon further inspection, we found that the number of duplicate clusters typically matched the number of *Ciona* adults used in each experiment.

**Figure 1:**
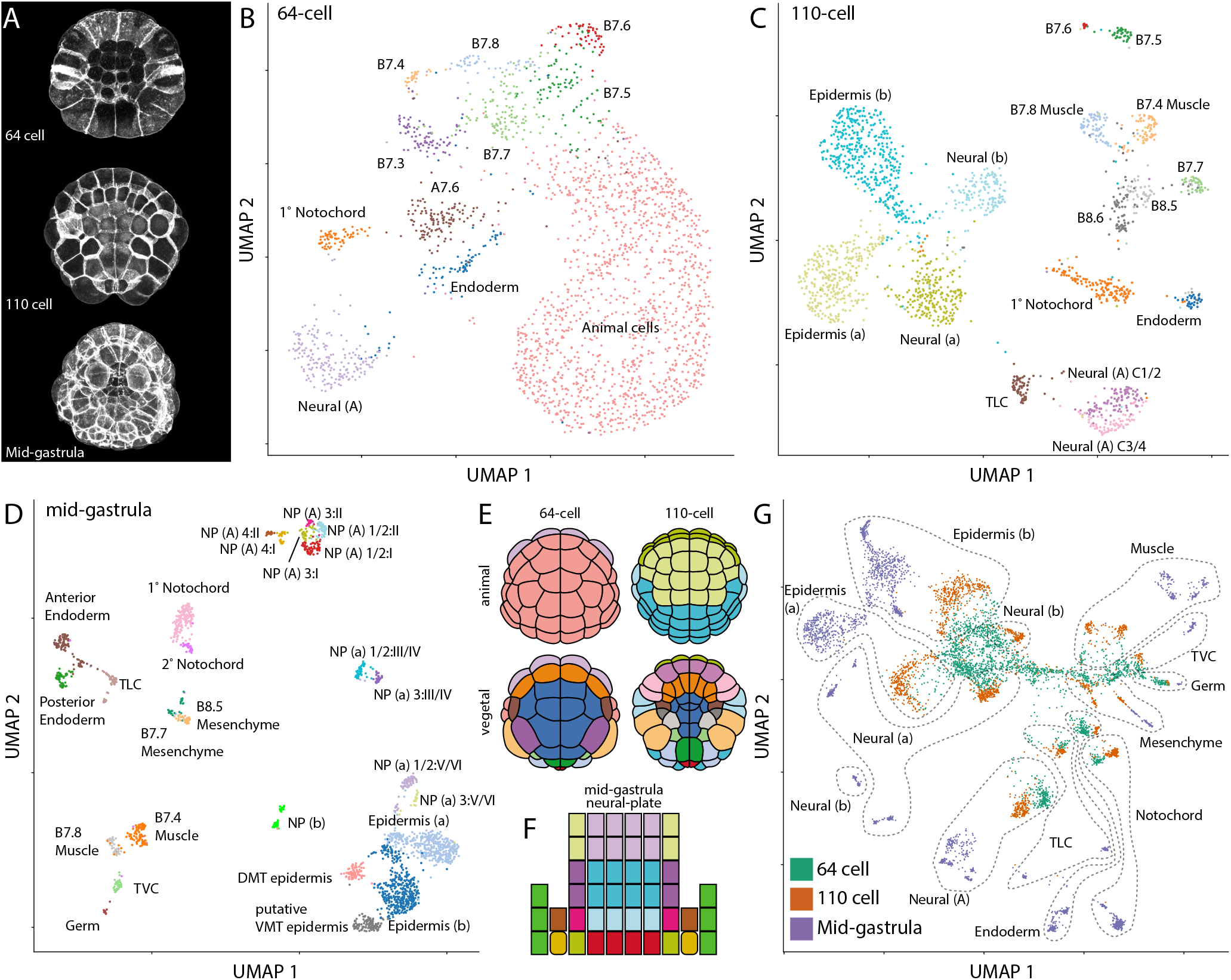
*Ciona* scRNAseq at the 64-cell, 110-cell and mid gastrula stages. A) Z-projections of confocal images of embryos at each stage analyzed. Anterior is up in all images. B-D) UMAP plot of STAMPs in control embryos collected immediately after the 64-cell (B), the 110-cell (C), and the mid-gastrula (D) stages. NP – Neural plate, NP X:Y represents the cell in row X and column Y of the neural plate. E) Cartoon representations of the animal and vegetal views of the embryo at the 64-cell and 110-cell stages. Colors correspond to the stage matched UMAP plot in B & C. F) Cartoon depiction of the neural plate at the mid-gastrula stage. Colors in F match the corresponding points in D. G) UMAP plot of all control STAMPs collected across all three developmental stages. Dashed lines delineate broad lineages of the embryo.

A recently published *Ciona* single-cell RNAseq study near the time of zygotic genome activation at the 16-cell stage found that single-cell transcriptome profiles tended to cluster not by cell type, but by embryo-of-origin [24]. As that study only profiled 4 hand-dissected embryos, each of which was from a different mother, they were unable to determine if these effects were due to differences between mothers of origin or individual embryos. To determine if differential deposition of maternal RNA between the different adults used in each experiment could be driving our observed duplication of cell type clusters, we took advantage of *Ciona’s* relatively high rate of polymorphism [49] and called SNPs across the entire genome for each STAMP in our data. We calculated a metric of relatedness between all pairwise comparisons of STAMPs [50] at each stage and hierarchically clustered the resulting relatedness matrix. The dendrograms obtained from clustering at each stage contained several long branches, which matched the number of adults used in each experiment (Figure S2 D-F). We then mapped the putative maternal identity back on the UMAP plots and found that the STAMPs tended to be clustering primarily by their mother at early stages, with this effect diminishing at later stages (Figure S2 G-I). This trend is consistent with expectations if differences in maternally deposited RNAs are contributing the majority of the variance between STAMPs early on and if this effect is “washed out” over time as zygotic transcription products accumulate in the cells. We then used the variable regression capability of the ScaleData function in Seurat [51,52], which creates a linear model for each gene expression feature in the data based on the effects of a given variable, to regress out the mother-of-origin effects and allow cells to cluster based strictly on cell type-specific gene expression.

Following post-sequencing SNP analysis and computational clustering, we again generated UMAP plots for each stage (Figure 1 B-D). This time we were able to assign cell identities to the scRNAseq clusters at all three stages, and the number of clusters was now in line with expectations based on extensive prior gene expression profiling (Figure 1 E-F and Supplemental Table 2). Some of these clusters represented broad territories of the early embryo such as the presumptive epidermis, but others had single blastomere-pair resolution such as the lateral columns of the posterior neural plate at the mid-gastrula stage. Expression patterns of the most highly variable transcription factors in the 110-cell stage clusters are generally similar to their previously characterized expression patterns by in situ hybridization but revealed important quantitative dynamics to expression patterns that had previously been appreciated only in a binary ON/OFF framework (Supplemental Table 3).

In addition to recapitulating almost all of the cell types expected at these stages based on the *Ciona* fate map and gene expression databases, we also identified three previously unappreciated transcriptional states in the mid-gastrula embryo: a cluster of putative posterior ventral midline epidermal precursors and two distinct cell states within the b-line neural lineage. The fate of all blastomeres in the 110-cell embryo have been previously analyzed [21], and it is known how the posterior animal blastomeres contribute to distinct dorsal-ventral and anterior-posterior locations in the tail epidermis at tailbud stages [35]. The ventral midline is known to be functionally and transcriptionally distinct from more lateral tail epidermis at later stages [35], but the ventral midline patterning mechanisms are not well understood. Here we identify a distinct cluster of posterior epidermal cells that we tentatively identify based on Nkx-A, Hox12, and Wnt5 expression [47,53] as the posterior-most cells contributing to the ventral midline tail epidermis at tailbud stages (descendants of b8.27 cell pair) (Figure 1 D). Additionally, there are two distinct clusters of b-line neural cells at the mid-gastrula stage that express distinct markers that have yet to be analyzed by in situ hybridization. Patterning of b-line neural cells is not well understood, so these new clusters provide an entry point for future studies.

### Linking cell states across time

*Ciona* embryos have stereotyped and intensively studied cell lineages, so the parent/daughter/sibling relationships between the cell states represented by different STAMP clusters are straightforward to infer. We first assigned cell types to each STAMP cluster separately at each stage. To confirm the consistency of these assignments between stages, we clustered the STAMPs from all three of our stages in the same reduction of high-dimensional gene expression space. The UMAP plot from this reduction represented the known lineages in *Ciona* embryos relatively well (Figure 1 G and Figure S1). The STAMPs from a given lineage across stages tended to cluster close together in a pattern that radiated outwards from the center of the UMAP plot with STAMPs that were later in developmental time being further from the center (Figure 1 G). This generally confirmed that our cell type assignments were consistent between lineages and showed an overarching trend that cell types established at the 64-cell stage or earlier tend to become far more transcriptionally divergent over time.

### Some FGF dependent cell types are not transfated to sibling cell types upon MAPK inhibition

To understand the transcriptional responses to FGF/MAPK signaling, we performed these scRNAseq experiments in parallel with control embryos and embryos treated with the MEK inhibitor U0126 from the 16-cell stage. Treatment with U0126 at this timepoint is known to block induction of mesenchyme, endoderm, and notochord [34,37,38]. It also prevents neural induction in two subsets of animal blastomeres [44] and disrupts the mediolateral patterning of the A-line neural plate, which secondarily prevents the establishment of A-line secondary muscle fate. The effects on A-line neural plate mediolateral patterning are mediated through a Nodal/Delta signaling relay downstream of FGF/MAPK signaling [42]. This Nodal/Delta relay is also required for the fate bifurcation event involving secondary notochord [43]. We found that most of the expected cell lineages are missing from the drug treated embryos (Figure 2 A) and that sibling cell types are present in excess. In order to define a metric of FGF dependence for each bifurcation, we predicted cell types in the U0126 STAMPs using the cell type label transfer function from Seurat. These predictions were inconsistent for certain cell types at the 64- and 110-cell stages but aligned closely with expectations from published literature at mid-gastrula as sibling cell types become more divergent. We performed chi-square tests for whether drug treatment influences the proportion of cells in sibling clusters (using the STAMPs for all descendant lineages of a STAMP cluster at mid-gastrula as a proxy) and took the negative log of the p-value as our metric. We were not able to compute this metric for the bifurcations where the parent cluster was itself FGF dependent. The only bifurcation that was not significant was the B7.5/B7.6 bifurcation which is believed to be FGF independent at this stage (Figure 2 B). For the U0126 sensitive bifurcations, there was substantial variability in the FGF sensitivity metric. This likely reflects a lack of sensitivity of the label transfer function we used to assign fates to the U0126 treated cells when the divergence distance in gene expression space of the FGF dependent cluster from their FGF independent siblings is small.

**Figure 2:**
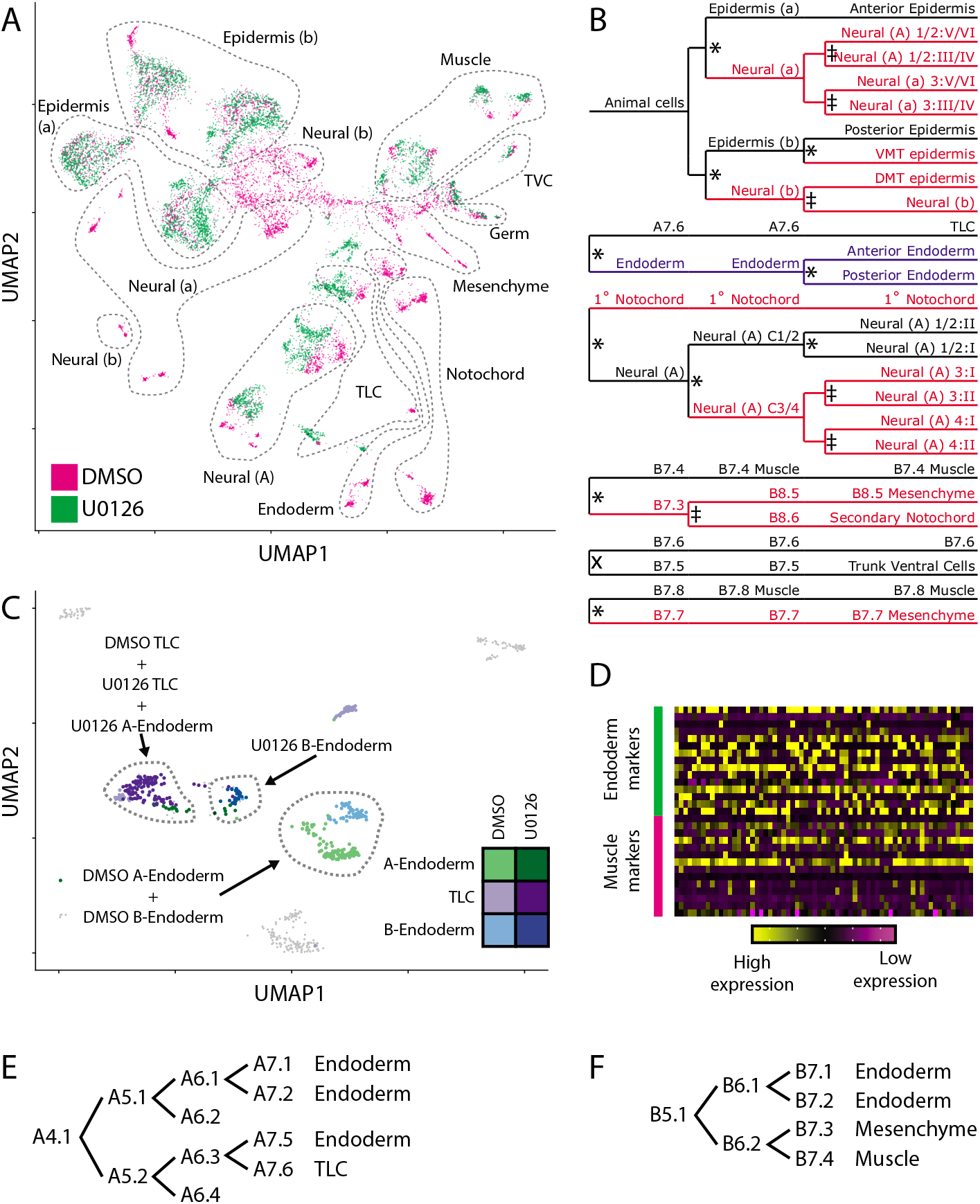
MAPK inhibition eliminates many cell types. A) UMAP plot of all DMSO and U0126 treated STAMPs across all three stages. UMAP space and dashed lines are identical to Figure 1G. B) Lineage diagram of all STAMP clusters identified at the 64-cell, 110-cell, and mid-gastrula stages. Red lineage branches are missing after treatment with U0126, black lineage branches remain after U0126 treatment, blue lineages branches are transfated to a unique state. *=p< 0.05, X=p > 0.05, ‡=untestable. C) UMAP plot of endoderm and TLC STAMPs at the mid-gastrula stages depicting the incomplete transfating of posterior endoderm. D) Expression of top endoderm and muscle markers in the U0126 treated posterior endoderm. E) lineage diagram of A-line endoderm from the 8-cell stage to the 64-cell stage. F) lineage diagram of the B-line endoderm from the 16-cell stage to the 64-cell stage.

All of the FGF-dependent bifurcations except one were consistent with the FGF-dependent cell type being transfated to its sibling cell type. We detected one novel cluster at the mid-gastrula stage that was only made up of STAMPs from U0126 treated embryos and not control STAMPs (Figure 2C). This cluster expresses some but not all markers of both endoderm and muscle and is thus likely to represent the B-line posterior endoderm (Figure 2D). Previous studies indicate that MAPK inhibition leads to A-line endoderm being transfated to A7.6 trunk lateral cell precursor fate (Figure 2E) [37] but have been unclear on what becomes of the posterior B-line endoderm. In a different tunicate, *Halocynthia roretzi,* FGF inhibition has been reported to cause the posterior B-line endoderm to adopt a cousin-cell muscle fate (Figure 2 F) in explants, but not in whole embryos [54]. We see the expected increase in STAMPS matching the A7.6 TLC precursor transcriptional profile, but this novel cluster indicates that the lineages that normally form B-line endoderm are not transfated to sibling cell types but instead establish a new transcriptional state not seen in normal embryos. Maternal β-catenin is a well-established endoderm determinant in ascidians and is involved in germ layer segregation early in development [29,30,32]. One interpretation is that vegetal FGF signaling acting downstream of maternal β-catenin is required for the normal posterior endoderm transcriptional regime, but that β-catenin alone is sufficient to make the B6.1 posterior endoderm lineage persistently distinct from the B6.2 muscle/mesenchyme/2° notochord lineage.

### Elk1/3/4 as a putative autoregulatory TF in an FGF dependent feedback loop

For all of the FGF dependent cell bifurcations, we wondered to what extent these diverse lineages exhibit a universal FGF/MAPK transcriptional response versus lineage specific responses. To address this, we clustered the fold-change in expression between FGF dependent lineages and their FGF independent siblings for the most variably expressed TFs. Most lineages exhibit their own characteristic responses, but we noticed a striking and unexpected pattern that the Ets family transcription factor Elk1/3/4 is consistently upregulated in the FGF dependent cell type compared to its sibling cell type (Figure 3 A). The FGF/RAS/MAPK/MEK signaling pathway culminates in *Ciona* and other animals in a transcriptional response largely mediated through Ets family transcription factors [55,56], of which the *Ciona* genome contains 13 putative members. Published in situ expression patterns of Elk1/3/4 at these stages are complex and hard to interpret, so it is possible that this relationship may have previously been missed without the quantitative data provided by scRNAseq. Elk1/3/4 expression is strongly MAPK dependent as demonstrated by its drastically reduced expression in U0126 treated embryos (Figure 3 B-C). We scanned for consensus Ets binding motifs in open chromatin regions throughout the *Ciona* genome using the early whole-embryo ATACseq data of [57] and found that the Elk1/3/4 locus is associated with a large amount of nearby open chromatin containing a very large number of predicted Ets sites (Figure 3 D-E). Elk1/3/4 expression is at least a proxy for Ets family transcriptional activity and given that Elk1/3/4 is itself an Ets family TF, it may have an important feedback role in FGF/MAPK-driven cell fate specification.

**Figure 3:**
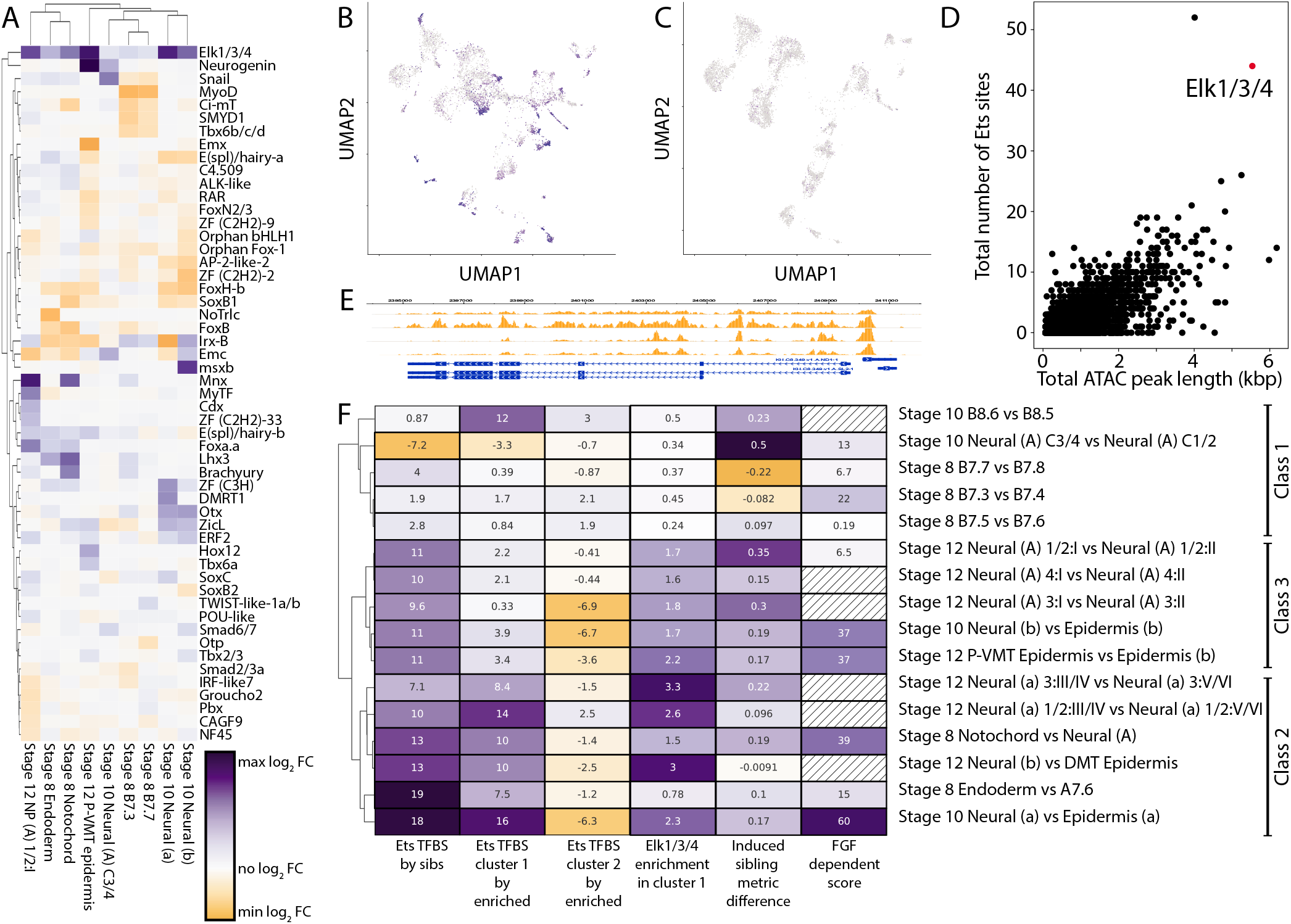
Elk1/3/4 is a putative feedback regulator of MAPK signaling. A) Log2 fold-change of the expression level of the top 50 most variably expressed transcription factors in U0126 sensitive cell type clusters compared to their U0126 insensitive sibling cell type clusters. B-C) Expression of Elk1/3/4 in DMSO (B) and U0126 (C) treated embryos. UMAP space is identical to figure 2A. D) Ets sites are enriched in the open chromatin regions near the Elk1/3/4 gene compared to all other gene models. E) Putative Elk1/3/4 sites in the 1500 bp upstream of the Elk1/3/4 transcription start site. The orange tracks show ATACseq open chromatin profiling at multiple stages from [57]. F) Ets TFBS and Elk1/3/4 expression enrichment in sibling cell types differentiate MAPK transcriptional response type. Ets TFBS by sibs – Ets TFBS score in the first sibling cluster compared to the second sibling cluster when using the set of genes upregulated in the first sibling compared to the second sibling for TFBS analysis. Ets TFBS by enriched – Ets TFBS score for that sibling cluster when using the set of genes upregulated in that cluster compared to all other clusters at a given stages for TFBS analysis. Elk1/3/4 enrichment – Difference in average expression level in sibling cluster 1 compared to sibling cluster 2. Induced sibling metric – Difference in total TF fold-change upregulation in sibling 1 compared to sibling 2. FGF dependent score – negative-log10(p-value) of the chi-square test for association between sibling cluster numbers and treatment status, hatched boxes could not be measured due to loss of a precursor cell type.

### Inferring the transcriptional role of FGF signaling in MAPK dependent cell fate bifurcations

The genome-wide transcriptional responses to FGF/MAPK signaling have not previously been assessed for the cell fate bifurcations captured here. We sought to measure aspects of this response to make inferences about the directness of the transcriptional dependence of FGF/MAPK signaling in FGF/MAPK dependent lineage decisions. To quantify the contribution of Ets family and other transcription factors to these FGF-induced cell states, we performed transcription factor binding site (TFBS) enrichment analysis. TFBS enrichment analysis uses statistical models to determine if the binding site sequence for a particular transcription factor (TF) is overrepresented in a set of target DNA sequences compared to a set of control DNA sequences [58]. In *Ciona,* most enhancers and other cis-regulatory modules are thought to be relatively close to the transcription start site [59]. We used the 1.5 kB of DNA upstream of the transcription start site to test for enrichment of Ets sites in the putative regulatory regions of genes that are upregulated versus downregulated in FGF-dependent cell types as compared to their sibling cell types. We also performed TFBS enrichment analysis in parallel using open chromatin regions inferred from whole-embryo ATACseq data [57] instead of the 1.5kb upstream regions. Following these two analysis methods, we averaged the enrichment metric z-scores for each TFBS from each method to get a combined z-score of binding site enrichment in target sequences compared to control sequences.

Z-scores for the Ets family TFBS vary widely across the different FGF-dependent cell fate bifurcations, indicating that Ets-mediated transcription is likely to play a major role in some bifurcations and not in others. Divergent use of Ets sites between sibling cell types could potentially reflect upregulation of Ets-mediated targets in the induced cell type and/or repression of Ets-mediated targets in the uninduced sibling cell type. To address this, we again performed TFBS enrichment analysis, this time using the genes that are up- and downregulated in a cell type compared to all other cell types at a given stage to make up the target and control DNA sequence sets. This was performed for each bifurcation for both the FGF-dependent cell type and its default sibling cell type.

To identify distinct trends of Ets transcriptional responses downstream of FGF/MAPK signaling, we performed hierarchical clustering of the enrichment z-scores for the Ets family TFBS calculated in the three different ways discussed above (Figure 3F columns 1-3). We also included differential expression of Elk1/3/4 between sibling cell types in the clustering given that we found it to be a likely proxy for Ets family TF activity (Figure 3 F column 4). The cell fate bifurcations clustered into 3 distinct classes. The first class represents bifurcation events that show little enrichment for Ets family TFBSs in the set of genes that differentiate the two sibling cell types and also have little to no upregulation of Elk1/3/4 in the induced cell type (“Class 1”). The remaining bifurcations all have strong upregulation of Elk1/3/4 expression in one of the sibling cell types, and are enriched for Ets family TFBS in the set of genes that differentiate the two sibling cell types. These bifurcations can be divided into one class that has strong Ets site enrichment in the induced cell itself when comparing genes that are enriched vs depleted in that specific cell type (“Class 2”), and another class where there is little or no such enrichment (“Class 3”). Many of this third class instead show Ets site depletion in the uninduced sibling. (Figure 3 F).

Prior studies have examined the role of FGF and other signaling pathways in some but not all of these lineage decisions. For example, the induction of both primary notochord and a-line neural fates are both thought to be directly dependent on FGF signaling [33,36,38,44]. Both of these bifurcations show strong enrichment of Ets TFBSs in both the induced cell type versus its uninduced sibling cell type, as well as comparing upregulated versus downregulated genes in the induced cell type alone (Class 2 in Figure 3 F). Both of these also showed strong enrichment of Elk1/3/4 in the induced cell type. Conversely, the induction of both column 3/4 lateral A-line neural plate and secondary notochord fates are known to be indirectly induced via the Nodal/Delta relay [42,43], and here show minimal Ets site enrichment comparing upregulated genes in the induced versus sibling cell types as well as little Elk1/3/4 enrichment. These results with well-characterized cell fate bifurcations support the idea that Ets TFBS enrichment and Elk1/3/4 expression differences provide a meaningful readout of Ets-mediated transcriptional activity.

All three bifurcations involving the posterior most rows of cells in the neural plate neural plate are thought to directly rely on FGF signaling to promote row I (posterior-most row) fate and repress row II (second posterior-most row) fate [41]. In our data all these bifurcations show Ets site enrichment in the row I cell types compared to the row two cell types, but only mild Ets site enrichment in the row I cell types comparing upregulated and downregulated genes (Class 3 in Figure 3 F). We interpret this to mean that Ets mediated transcription plays a role in differentiating the row I and row II cell types but plays less of a role in shaping the overall transcriptional state of the row I cell types. These results are in line with the previous understanding of A-line neural gene regulatory networks, where FGF activity in row I is known to control differences in transcription between rows I and II but is not thought to play a critical role in defining the A-line neural lineage overall.

The group of bifurcations that show little to no Ets TFBS enrichment (Class 1 in Figure 3 F) contains the expected bifurcations involving secondary notochord versus mesenchyme and the medial versus lateral A-neural lineages, which are known to be indirectly downstream of FGF [42,43]. This group also contains the bifurcation between the TVC precursor and the germ line precursor cell, which is known to be dependent on the CAB complex and not on FGF signaling [60]. Unexpectedly, however, this group also contains the bifurcations that differentiate the muscle lineages from their sibling cell types, including mesenchyme and 2° notochord, at the 64-cell stage. These bifurcations are thought to be directly dependent on FGF [34] but do not show an obvious transcriptional signature of Ets-mediated transcription in our analysis. These are unusual divisions in that FGF signaling is here thought to act by repressing in some way the activity of a maternal determinant of muscle fate [34]. This indicates that the FGF signal necessary for mesenchymal fate in these mesenchyme/muscle bifurcations could either operate indirectly of canonical Ets family TF-mediated transcription, or else it could lead to changes in the expression of only a few key genes downstream of Ets family members.

TFBS enrichment analysis of the transcriptome-wide changes in sibling cell lineages after fate bifurcation implies that the majority of fate bifurcations at these stages rely at least in part on direct input of Ets family TFs, presumably downstream of FGF signals. This includes the novel bifurcation that establishes the putative ventral midline tail epidermis at mid-gastrula. Other than the medial vs lateral A-line neural division that is well established as being indirectly downstream of FGF via the Nodal/Delta relay, the other 4 bifurcations with minimal signs of Ets-mediated transcription all involve posterior vegetal B-line cells. This includes one bifurcation expected to be indirect (B8.6 vs B8.5), one bifurcation that is not FGF dependent at all (B7.6 versus B7.5) and the two bifurcations involving primary muscle fate where the lack of a major Ets enrichment signature was unexpected.

### SMAD mediated repression of Ets site transcriptional targets may function in medio-lateral patterning of the neural plate

Nodal signaling is transcriptionally mediated through SMAD family TFs. Previous research shows that Nodal signals are responsible for the mediolateral patterning of the *Ciona* neural plate [42]. Interestingly however, one of the strongest TFBS enrichment signatures in the bifurcation between lateral and medial A-line neural siblings (column 1/2 vs column 3/4 at the 110-cell stage) is not an enrichment of SMAD family TF sites in the lateral A-line neural plate siblings, but rather a depletion of Ets family TF sites in the lateral neural plate siblings (Supplemental Table 4). There is precedent in other *Ciona* lineages that SMADs acting downstream of TGF-β superfamily signaling molecules can act to repress transcription of Ets family TF targets [61]. This implies that the Nodal signal that serves to pattern the medial-lateral axis of the A-line neural plate may function at least in part to repress Ets site enriched target genes in lateral cells and not strictly through the induction of directly SMAD-dependent targets.

### Analyzing the genome-scale dynamics of cell fate induction

Our dataset’s wide coverage of cell types and stages enables a systematic analysis of the transcriptional changes underlying cell fate bifurcations. We first explored the expression changes of transcription factors in “trios” of STAMP clusters involving a mother cell type and its two daughter/sibling cell types. For transcription factors that passed a statistical test for differential expression between the two sibling cell clusters, we compared the expression level of that transcription factor in all three members of the trio (Figure 4 A and Supplemental table 5). These heatmaps reveal several consistent features. Numerous transcription factors are differentially expressed between the sibling cell states in each trio comparison. The majority of these were already expressed to some extent in the parental cell type, but some are newly expressed in one or both daughter/sibling clusters. One daughter typically has more upregulated TFs than the other (Figure 4 A).

**Figure 4:**
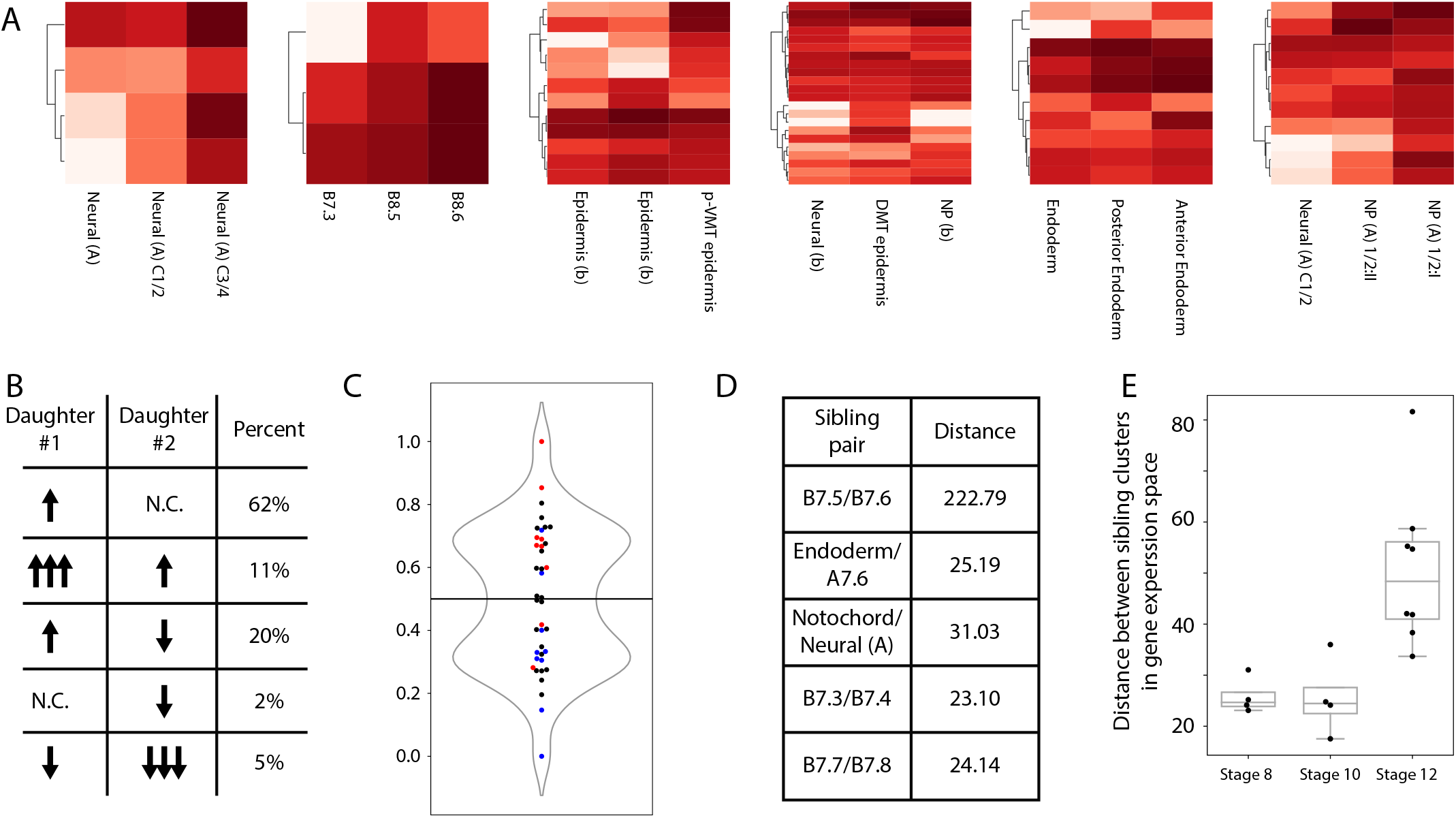
Genome-wide transcriptional responses during fate bifurcation events. A) Log2 expression level of differentially expressed transcription factors in “trios”. For each heatmap, the left column represents the parent cell type, the middle column is the “uninduced” sibling cell type, and the right column is the “induced” cell type (See also supplemental table 5). B) Different mechanisms of changes in expression level that lead to upregulation in daughter #1 compared to daughter #2, and percentages observed of each mechanism in our “trios”. C) A simple “induction metric” can be used to predict FGF sensitivity. Blue = U0126 insensitive cell types by chi-square test, Red = U0126 sensitive cell types by chi-square test, black = untestable cell types by chi-square test (corresponds to figure 2B). D) Distances between sibling cell types in gene expression space for all bifurcations occurring at the 64-cell stage. E) Distances between sibling cell types in gene expression space for all bifurcations occurring at each stage (Germ lineage not included).

If a transcript is upregulated in cell type A compared to its sibling cell type B, there are several possible underlying mechanisms (Figure 4 B). It could either be (1) upregulated in daughter A vs the parental cell state and unchanged in daughter B vs the parental state, (2) upregulated in daughter A vs the parental state and upregulated in daughter B vs the parental state, but upregulated more in A than in B, (3) upregulated in daughter A vs the parental state and downregulated in daughter B vs the parental state (4) unchanged in daughter A vs the parental state and downregulated in B vs the parental state, or (5) downregulated in both daughters versus the parental state, but downregulated more in daughter B than in daughter A.

We found that 62% of TFs that are differentially expressed between sibling cell types in a trio fall into the first category, 11% fall into the second, 20% into the third, 2% into the fourth, and the remaining 5% in the fifth category. (Figure 4 B) The largest driver of differential expression between sibling lineages is upregulation in one sibling and not the other, but a surprising fraction of genes were upregulated in both siblings but more so in one than the other.

The trio comparisons showed that one daughter typically had more upregulated transcription factors than the other, and we found that this was also true in the other bifurcations where we did not capture the parental cell state (Supplemental File 1). We hypothesized that this might reflect which daughter cluster in a cell fate bifurcation was induced downstream of FGF signaling. We tested this by calculating a simple metric of upregulated TF expression between sibling cell types, by determining what proportion of the total upregulation of differentially expressed TFs was in each sibling cell type. This putative induction metric correctly predicted the induced cluster in almost all cases (Figure 4 C and Figure 3 F). The only exceptions were the B7.4 and B7.8 muscle lineages, which had more upregulated TFs than their FGF-dependent mesenchyme and mesenchyme/2° notochord precursor sibling cell types. Muscle cell fates are known to be driven, however, by asymmetric segregation of a cell-fate determinant (macho-1/Zic-r.a) [62], and these cells could be considered “intrinsically induced”.

### Newly born cell fates diverge from their sibling fates more quickly at later stages

The distances in gene expression space between sibling cell fates after bifurcation are widely variable across developmental time and anatomical regions. Some sibling cell types remain fairly close, indicating that gene expression profiles are not drastically changed between sibling cell states, whereas others are more divergent. The relatively transcriptionally silent B7.6 cell and its sibling blastomere B7.5 are far more distant from each other than other sibling cell state pairs at the same stage (Figure 4 D). This likely reflects maternal cell fate determinants known to segregate as RNA molecules into the B7.6 germ cell precursor downstream of patterning by the CAB [63]. We do however find a consistent trend that at the mid-gastrula stage, the distance between sibling STAMP clusters which have just bifurcated is much larger than the distance between sibling STAMP clusters immediately after bifurcation that split at the 64-cell or 110-cell stage (Figure 4 E). This trend is also readily visible by looking at the distance between newly born sibling cluster in UMAPs space at each of the three stages (Figure 1 G). This indicates that the rate of divergence in gene expression space between newly established sibling clusters is increasing with developmental time, and potentially reflects the establishment of new chromatin states and/or the zygotic expression of a broader set of transcription factors as development proceeds.

### Many cell fate bifurcations involve large numbers of differentially expressed transcription factors

Exploring the different cell fate bifurcations, we found that most involved a surprising number of differentially expressed TFs. Supplemental file 1 is a large poster-sized diagram detailing all of the differentially expressed transcription factors across all of the bifurcations analyzed. Decades of work in developmental genetics have emphasized the role of single transcription factors or small combinations of transcription factors in controlling specific cell fate decisions, but we found that many bifurcations involved 10 or more different TFs becoming differentially expressed between sibling cell types within an hour or less of cell division. This led us to explore the dynamics of these transcription factors over time to identify if they remained expressed in the tissue throughout development, or if they were only transiently upregulated at the time of differentiation. To address this, we used our scRNAseq dataset to identify all of the transcription factors upregulated in the earliest stages of muscle, mesenchyme and endoderm differentiation as compared to their sibling cell types. We selected these cell types because they formed entire tissues at later stages of development without the dramatic increase in tissue subtypes exhibited in the neural lineages. Notochord is discussed in more detail in a later section but shows similar trends. Each of these tissues exhibited differential expression of a suite of several transcription factors. (Figure 5 A-C). To infer patterns of transient versus stable expression, and to explore more of the temporal expression dynamics of these earliest diverging TFs, we also integrated our scRNAseq dataset of early *Ciona robusta* development with another single-cell RNAseq dataset focused on later *Ciona robusta* stages [10]. We compared the expression level from 64-cell through hatching larva of the TFs which were upregulated in muscle, mesenchyme and endoderm compared to their sibling cell types at their initial bifurcation. We find that these TFs generally fall into one of two categories: some are expressed for only a short time window whereas others show persistent and/or increasing expression (Figure 5 A-C). Functional experiments are needed to determine which transcription factors diverging between sibling cell types have important roles in cell fate specification, but all of the bifurcations we have examined show several transcription factors becoming differentially expressed instead of a single putative master regulator. All of these newly-specified cell types were captured by scRNAseq very soon after being separated from sibling lineages by cell division. They were typically on ice during the dissociation process within 15 minutes of the prior division and microfluidic encapsulation and lysis were underway within 35-40 minutes. It is thus likely that these relatively large suites of divergently expressed TFs represent direct responses to the earliest aspects of cell fate bifurcation.

**Figure 5:**
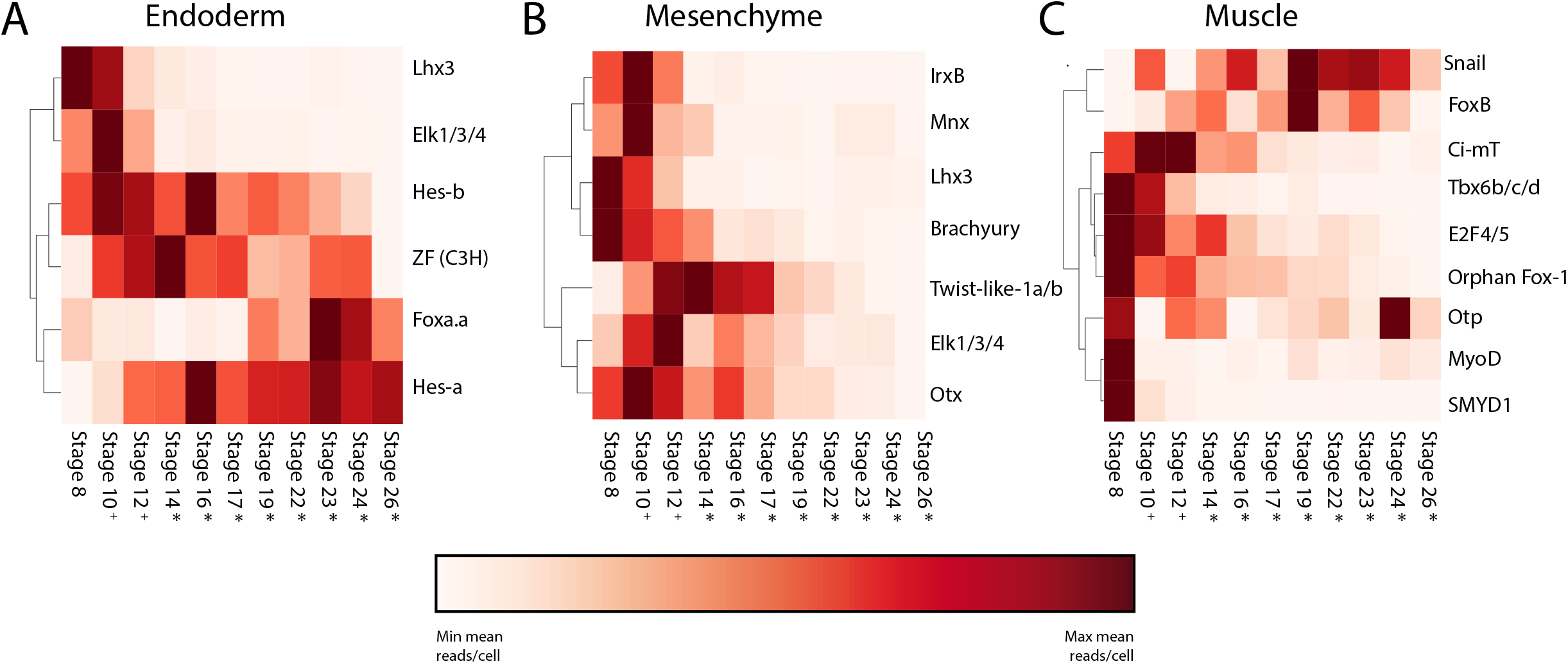
Differential transcription factor expression in cell fate bifurcations. A-C) The expression level over time of TFs that are differentially upregulated in the Endoderm (A), Mesenchyme (B), and Muscle (C) lineages compared to their sibling cell types.

### Many genes besides *Brachyury* are differentially expressed in the notochord immediately after fate bifurcation

The *Ciona* notochord has been intensively studied as a model for understanding tissue-specific gene expression and morphogenesis [64–73]. This is the first study to systematically identify DE genes in the *Ciona robusta* primary notochord compared to its sibling A-line neural cell fate immediately after cell fate induction at the 64-cell stage. To analyze if these genes were in fact notochord specific, we extracted the average expression value of each of these genes in all of the other cell clusters at the 64-cell stage. We found that nearly all of the genes are expressed most strongly in the notochord but many are also expressed elsewhere in the embryo to varying extents, indicating that they are notochord enriched, but not notochord specific (Figure 6 A). There are 12 transcription factors and many other genes that diverge between the 1 ° notochord and its A-neural siblings at this first timepoint immediately after they divided. Many of these genes have not previously been identified as notochord enriched at the 64-cell stage and may be effector genes of the earliest stages of the notochord GRN. There is very little overlap between the genes differentially expressed in the notochord at the 64-cell stage and those induced by ectopic expression of Brachyury in other studies (Figure 6 B) [74,75], suggesting that these earliest differentially expressed genes may be expressed in parallel to, and not downstream of Brachyury. The transcription factors we found to be enriched in primary notochord as compared to A-line neural included several expected candidates such as Brachyury, Foxa.a and Mnx [47,48,72,76] as well as others such as Zic, Elk, and Hesb that have not been previously appreciated by *in situ* hybridization at the 64-cell stage. Much like in the endoderm, muscle, and mesenchyme, we find that the DE TFs are either expressed strongly in a short time frame around fate induction or are stably expressed and increase their expression over time (Figure 6 C).

**Figure 6:**
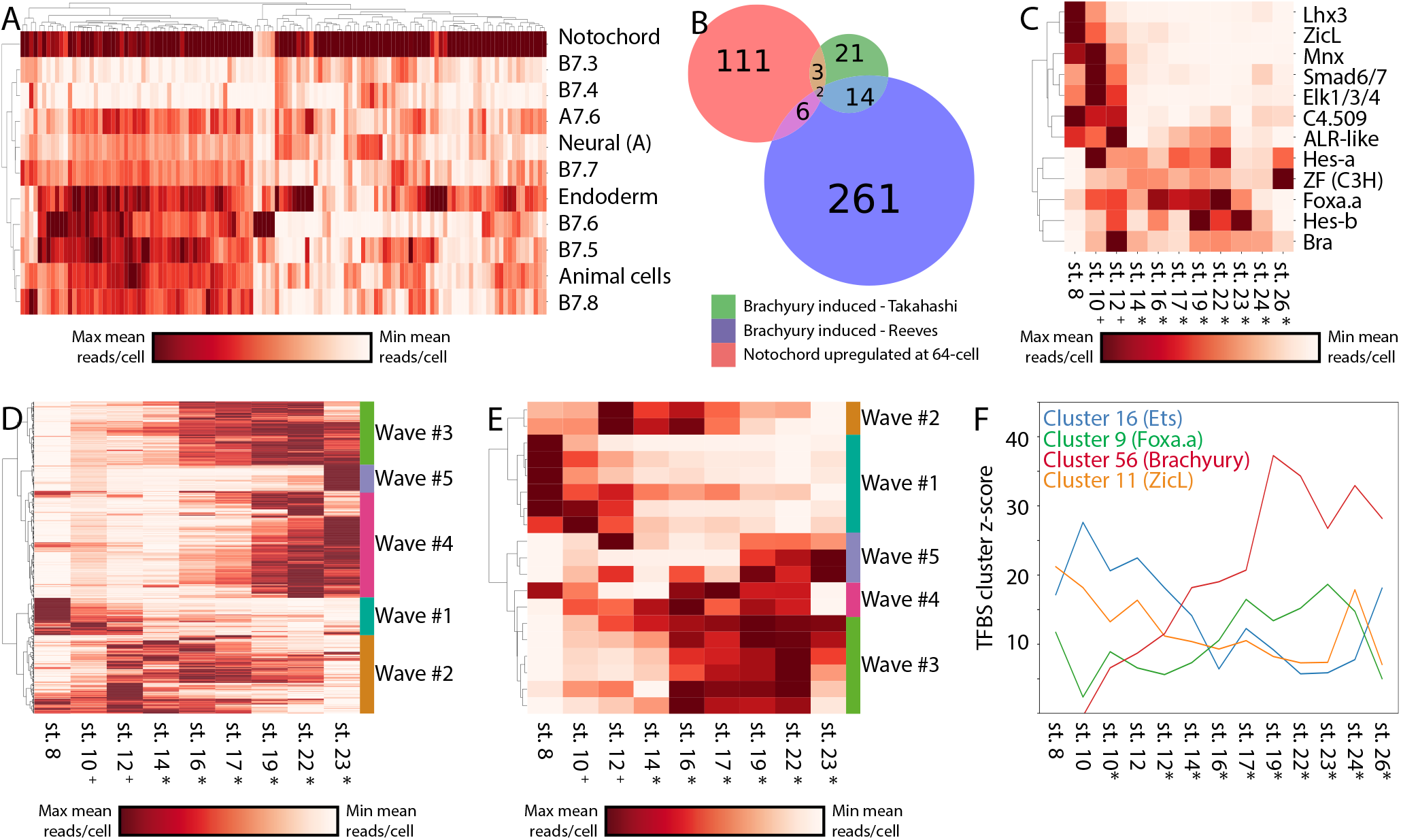
Waves of gene expression in the primary notochord. A) Spatial expression profiles of all genes that are upregulated in the 1 ° notochord compared to their A-neural siblings at the 64-cell stage. B) Overlap between the 1 ° notochord differentially upregulated gene set and two sets of Brachyury induced notochord-enriched genes from the indicated studies. C) Temporal expression profiles of TFs that are notochord upregulated at the 64-cell stage compared to their A-neural siblings. D-E) Temporal expression profiles of all notochord enriched genes (D), and transcription factors (E) reveal distinct waves of expression throughout the course of development. F) Enrichment z-scores for several TFBS clusters during the course of notochord development.

### The notochord exhibits distinct waves of transcription

To further characterize the temporal dynamics of the *Ciona* notochord transcriptome, we plotted the notochord expression levels over time of all the genes that are enriched in the notochord compared to other non-notochord tissues at each stage. Hierarchical clustering of these notochord enriched genes revealed 5 relatively distinct temporal waves of expression (Figure 6 D-E). Characteristic temporal expression profiles have previously been identified by in situ hybridization for select notochord genes, [66–68,71,72,74,77,78] but the genome-wide transcriptional profiling performed here allows the comprehensive identification of distinct suites of genes that co-vary in expression in the notochord over time. A similar set of distinct temporal waves was seen when clustering the temporal expression profiles of the notochord-enriched transcription factors alone, suggesting that these waves might be controlled by distinct TFs or combinations of TFs acting in a differentiation cascade. All the waves of TF expression in the notochord involve suites of multiple co-varying TFs and not single genes. This suggests that the temporal dynamics of notochord-specific gene expression might represent a progression through a series of relatively distinct transcriptional states that each involve TFs acting in complex sets and not as single regulators. The secondary notochord also exhibits distinct transcriptional waves involving an overlapping but non-identical set of genes. (Figure S3 A-B). Future work will be needed to dissect these new GRN structures and test whether these waves of transcription coincide with, and possibly contribute to the control of distinct morphogenetic events such as mediolateral intercalation and notochord elongation.

### The earliest notochord GRN is enriched for targets of Ets and Zic transcription factors

We used transcription factor binding site enrichment to infer potential roles for the numerous TFs enriched in the 1 ° notochord compared to its sibling cell type at the time of notochord induction. Using our scRNAseq dataset integrated with the partially overlapping Cao et al dataset [10], we performed TFBS enrichment analysis to identify which TF family binding sites are most enriched in the putative regulatory regions of genes enriched in primary notochord compared to other cell types at each stage. Our assumption from the current understanding of the notochord GRN is that the key notochord regulator Brachyury is induced by the combination of active Ets, Zic and Foxa.a, and that most notochord-specific transcription should be directly or indirectly downstream of Brachyury. Interestingly, we find that the earliest targets are not enriched for Brachyury sites, but are instead highly enriched for the binding motifs of Zic and Ets family TFs (Figure 6 F and Figure S4 A). Brachyury sites do not become enriched in notochord-enriched genes until much later in development, suggesting that there is a significant lag time between when it is first detectable at the RNA level by in situ at the 64-cell stage and when it becomes a major driver of notochord gene expression (Figure 6 F). This supports the hypothesis that most early notochord genes are initially induced directly downstream of FGF, in parallel to and not downstream of the key notochord regulator Brachyury [74]. We also see that Foxa.a has a U-shaped TFBS enrichment profile over time (Figure 6 F) and is one of the most enriched TFBS motifs in the later stages of development (Figure S4 B), consistent with a role in a recently described feedforward network as both an upstream regulator of Bra expression and a major regulator of notochord-specific gene expression at later stages [74].

## Conclusions

### scRNAseq in an outbred, polymorphic species

Whole-embryo scRNAseq is a powerful new method for generating transcriptional atlases of distinct cell states and understanding the transitions between them. Here we found that a modest number of whole-embryo scRNAseq experiments has the power to recapitulate many, though by no means all, key findings from years of classical developmental biology experiments. We were not able to detect the b-line neural cells as a distinct cluster until the 110-cell stage, but we otherwise detected all of the cell types expected from this simple and intensively studied embryo. scRNAseq also allowed us to uncover previously undescribed transcriptional states during early *Ciona* development that will facilitate studies of later b-line neural patterning as well as the patterning of the future tail epidermis.

An unexpected complication in our dataset was the tendency of some cells to cluster at early stages based on mother of origin and not strictly by cell type. We introduced a method for removing these effects post hoc by inferring the mother-of-origin for each STAMP using SNPs and regressing out the maternal influence. This method exploits the extensive natural genetic variation in *Ciona* populations, and will likely be useful for future scRNAseq studies in *Ciona* and other outbred, highly polymorphic species. SNP analysis could potentially be integrated with scRNAseq in a fecund, polymorphic species like *Ciona* in many interesting ways. STAMPS could potentially be assigned to distinct embryos to dissect variability on a per embryo basis. scRNAseq capture rates could be increased by overloading encapsulation devices and computationally identifying and removing cell doublets post hoc. This would be akin to the demuxlet algorithm currently used for multiplexed samples from different genotyped mouse or human samples [79] but now exploiting the much greater polymorphism of *Ciona* to infer post hoc genotypes for cells from thousands of different embryos resulting from mixing gametes from several hermaphroditic adults. Naturally occurring polymorphisms could also potentially be linked to transcriptional differences between genotypes.

### Global patterns of divergence between sibling cell types

*Ciona* and other tunicates are unusual among the chordates in having a modest repertoire of distinct cell types that become established via exceptionally stereotyped embryonic cell lineages. This makes it possible to achieve relatively deep coverage of most cell types in whole embryo scRNAseq experiments using standard commercialized droplet microfluidics, and also to capture newly divergent sibling cell types within minutes of having divided from common parental cell types. Our goal here was not just to create a transcriptional atlas of cell states during the late cleavage and early gastrula stages when most cells become_restricted to a single predominant fate, but also to systematically analyze the genome-wide transcriptional changes in the cell fate bifurcations that establish these distinct cell types.

*Ciona* development is unusual in that a large number of cell fate decisions are controlled by only a few signaling pathways, with a particularly important role for FGF signaling on the vegetal side of the early embryo [18,80]. A noteworthy feature of the cell fate bifurcations we profiled is that they were typically asymmetric in the sense that one sibling cell state had a greater number of upregulated transcription factors than the other. In almost all cases, the cell type with the greater number of upregulated TFs was the FGF-dependent cell type and not its FGF-independent sibling. This suggests that these inductive interactions are dominated by the upregulation of FGF target genes in the induced cell type. Interestingly, however, in the cases where we profiled parent-sib-sib cell state trios, the majority of TFs differentially expressed between the sibling cell types were already expressed to at least some extent in the parental cell type, indicating that the de novo expression of regulators unique to the induced cell state is not the dominant pattern.

Transcripts can become differentially expressed between sibling cell types through varying combinations of upregulation and downregulation in the two lineages. We found that the most common pattern was for a gene to be upregulated with respect to the parental state in one cell type while not changing in the sibling. All possible combinations were observed, however, and there were a surprising number of differentially expressed genes that were upregulated in *both* sibling cell types as compared to the parental cell type, but to varying extents. Note that differential transcript levels between sibling cell states might represent different underlying combinations of changes in transcription rates, decay rates, and asymmetric inheritance in cell division. The observation of strong TFBS enrichment signatures in many bifurcations supports the obvious role of direct transcriptional upregulation, but differential degradation and asymmetric inheritance could also be important in these very fast-paced cell state transitions.

Another noteworthy feature of the 17 bifurcations we examined is that later bifurcations were far more transcriptionally divergent than early bifurcations. We speculate that this might involve chromatin states becoming more open over time, and/or a larger set of zygotic transcription factors being expressed in parental cell states at later stages.

### *Ciona* superenhancers?

We found that the Ets family transcription factor Elk1/3/4 was unique in being upregulated in almost all of the MAPK-dependent cell fate bifurcations, especially in the ones thought to be directly dependent on FGF signaling and not downstream relay mechanisms. Vertebrate Ets proteins generally act as transcriptional activators, but repressive functions have also been described [81]. It seems likely that *Ciona* Elk1/3/4 is acting in a feedback loop, though it remains to be determined whether it is a positive or a negative loop.

*Ciona elk1/3/4* is distinctive in being associated with more extensive regions of nearby open chromatin than all but two other *Ciona* genes, and also with a very high number of predicted Ets sites in these putative regulatory regions. Studies of mammalian genomes have led to the hypothesis that key cell fate specification genes are often under the cis-regulatory control of unusually large regulatory regions known as super-enhancers [82–84]. The super-enhancer concept has not previously been applied to studies of the small, compact *Ciona* genome, where many genes are known to be regulated by very short enhancer sequences [57,59], but *elk1/3/4* suggests that the concept may be relevant. ChIP-seq data would be needed to formally test the mammalian super-enhancer criteria, but we find that transcription factors are enriched in the genes with more than 2kb of associated open chromatin (supplemental table 6), and that the transcription factor loci with the most associated open chromatin include the effector TFs of many signal transduction pathways with roles in early *Ciona* patterning. These include Elk1/3/4 and another Ets factor, RBPJ, a Lef/TCF and a Smad (supplemental table 7).

### The ‘Broad-Hourglass’ model of cell fate bifurcation

The gene regulatory networks controlling cell fate specification events are conventionally thought of as hourglasses with narrow waists (Figure 7A). Upstream of the cell fate specification event, diverse regulatory interactions give rise to some unique combination of lineage-specific transcription factors and signaling states that induces the expression of either a single master regulatory transcription factor unique to that cell type or a small number of tissue-specific transcription factors that define a combinatorial code. Transcriptional cascades acting downstream of that single factor or small set of factors then lead directly or indirectly to changes in the expression of large numbers of downstream genes. There are major questions, however, about how narrow the pinch point of the GRN hourglass actually is as sibling cell states diverge. Here we found that these bifurcations were transcriptomically complex, with many genes, including numerous transcription factors, differentially expressed between sibling cell states within minutes of cell division (Figure 7 B). We refer to this as the ‘broad hourglass’ to distinguish it from the ‘narrow hourglass’ model in which the earliest transcriptional divergence between sibling cell types involves no more than a few key transcriptional regulators of the newly-induced cell state. The hourglass metaphor is an apt way to think about the establishment of new transcriptional states, but note that we are not making reference to the concept of the phylotypic stage where an hourglass metaphor has also been widely used [85].

**Figure 7:**
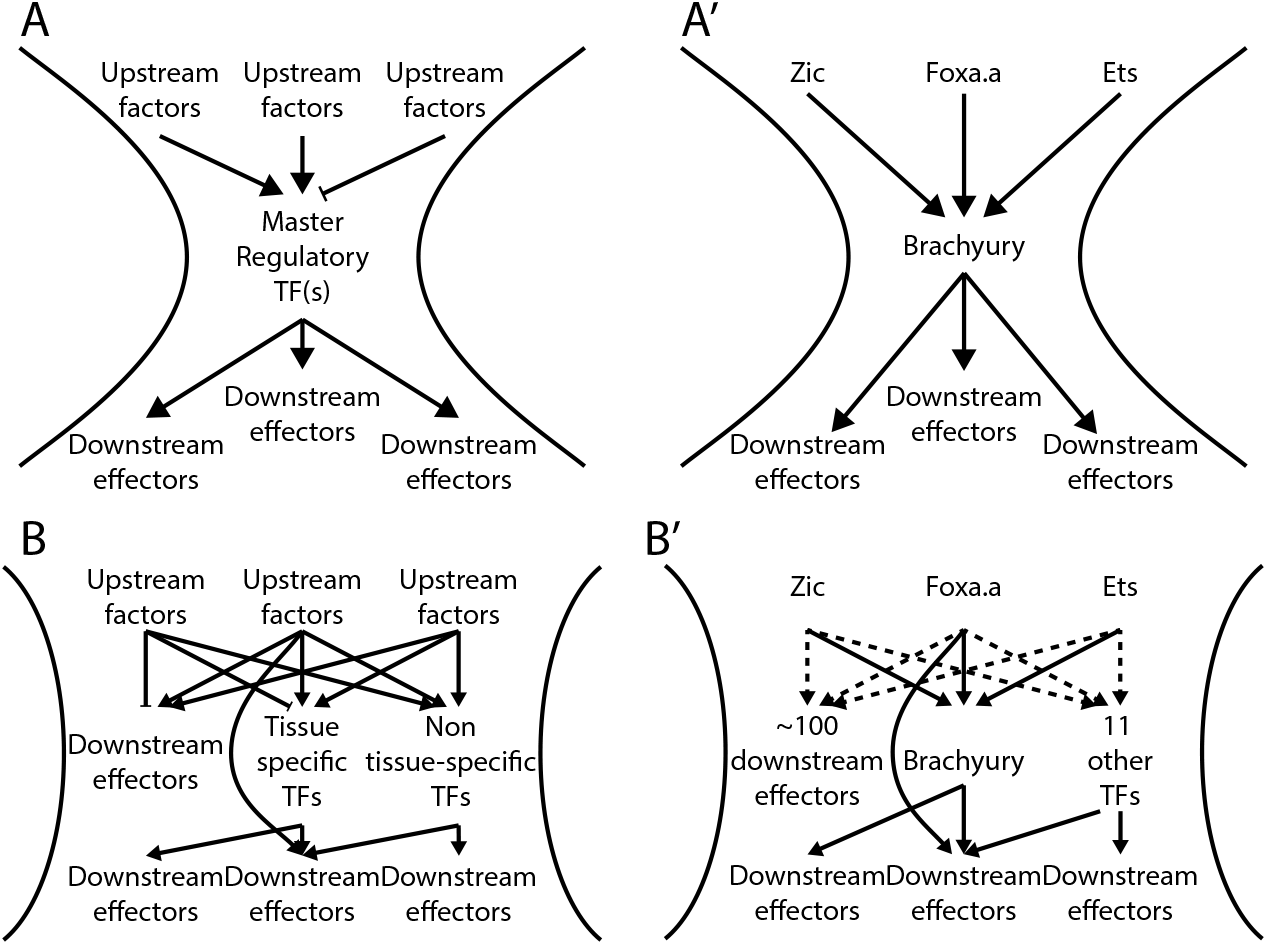
The wide-hourglass model of cell state transitions. A) A theoretical framework of a ‘narrow-hourglass’ transcriptional response being primarily deployed through induction of a single tissue-specific master regulator. A’) A narrow-hourglass model of the *Ciona* notochord GRN focused on Brachyury. B) The wide-hourglass transcriptional response includes many different TFs being upregulated alongside numerous putative effector genes at the time of cell fate induction via the same set of transcriptional inputs. B’) The wide-hourglass model of the *Ciona* notochord GRN presented in this paper. Dashed lines indicate transcriptional inputs broadly inferred from TFBS enrichment data, but these may not apply to each and every transcript upregulated in the early notochord.

The number of differentially expressed TFs observed across many different cell fate bifurcations raises interesting questions about what fraction of these TFs are functionally important for the bifurcations in which they become differentially expressed. Forward and reverse developmental genetic approaches have generally emphasized the necessity of individual transcription factors for specific cell fate decisions, but that does not exclude more complex sets of TFs playing important roles. Across many different fate bifurcations, we found that the earliest DE TFs consistently fell into two major classes. Some were stably enriched in the new cell type, whereas others were only transiently enriched. It remains to be determined to what extent transiently enriched TFs represent separable roles in the induction but not maintenance of cell fate as opposed to purely ephemeral responses.

In the bifurcation between primary notochord and A-line neural cell fates, we identified 12 differentially expressed TFs at the 64-cell stage. Some of these, such as Bra and Foxa.a, have well-established roles in notochord fate. Bra was until recently thought to be a master regulator of notochord fate, but it is increasingly clear that Foxa.a has important roles within the fate-restricted notochord lineage and not just as an upstream regulator of Bra expression [72,74]. We also found that Zic becomes enriched in primary notochord compared to its A-line neural sibling cells. Zic is similar to Foxa.a in having a well-established role as a direct upstream activator of Bra expressed at the 32-cell stage in the parent blastomeres of notochord and A-line neural cells [86,87] but has not previously been found to become differentially expressed between these cell types. Two other notable TFs that are enriched in primary notochord at the 64-cell stage include Elk1/3/4, which appears to be broadly involved in FGF/Ets mediated cell fate bifurcations, and also Hesb, which commonly acts as a transcriptional target of Delta/Notch signaling. Delta/Notch signaling is thought to be involved in inducing secondary but not primary notochord fate [43], but the lateral-most primary notochord cells are in contact with Delta2 expressing cells and Hes expression in these cells has been observed by in situ hybridization [43].

In addition to finding many transcription factors other than Bra and Foxa.a enriched in the presumptive primary notochord, we were also surprised to detect Bra expression in certain lineages outside the notochord. The secondary notochord lineage first becomes fate-restricted to notochord in B8.6, but we saw considerable Bra expression in its parent, B7.3, and also to a smaller degree in its sibling B8.5. Bra expression in B7.3 is seen in some but not all published in situ patterns, so this was not entirely unexpected. It may represent an example of mixed lineage transcriptional priming. Much more unexpectedly, however, we detected modest but statistically significant Brachyury upregulation in the row I column 3 A-line neural plate cells at the mid-gastrula stage compared to their row II sibling cell fate. These cells are similar to notochord in that they express Foxa.a, Zic and have high MAPK activity [41,56,88]. In vertebrate embryos, Brachyury has complex roles in posterior neuromesodermal precursors distinct from its role in the notochord [89–91]. Brachyury mutant embryos do not show an obvious neural tube defect [92], but we speculate that this expression domain might represent a vestigial remnant of an ancestral role in tail neuromesodermal lineages. This group of cells is fate-restricted to lateral tail nerve cord, but the A6.2/A6.4 lineage overall gives rise to a complex mixture of neural and mesodermal fates. Alternatively, Bra expression in this subregion of the neural plate might reflect partially overlapping cis-regulatory inputs, with upstream factors beyond Foxa.a, Zic and active Ets controlling whether Bra expression is transient or persistent.

Regardless of whether they are all functionally important, it is clear that many genes and not just one or two master regulators become differentially expressed between newly established *Ciona* cell types. A conventional view of *Ciona* primary notochord fate specification would be that the intersection of active Ets, Zic and Foxa.a expression induces Bra, and that Bra then induces early notochord target genes (Figure 7 A’). Here we propose that the early notochord-specific transcriptional regime is instead regulated by the same combination of active Ets, Zic and Foxa.a expression that induces Bra, and is thus in parallel to and not downstream of Bra activity. In support of this, TFBS enrichment analysis shows that Zic and Ets binding motifs are strongly enriched in the putative regulatory regions of early notochord-enriched genes and it is not until later that Bra motifs become strongly enriched (Figure 6 F and Figure 7 B’). It is perhaps unsurprising that the combination of active upstream transcription factors that induces a particular cell type would upregulate more than just a few tissue-specific master regulators, but it is surprising that it is not until much later that Bra becomes the predominant TFBS enrichment signature in the fate-restricted notochord. Tissue-specific effector networks

In addition to the early embryonic gene regulatory networks that establish distinct cell types, there are also gene regulatory networks active within distinct cell types that control the temporal aspects of gene expression that help drive the differentiation and morphogenesis of different tissues. These tissue-specific effector networks have received less attention than the early networks that initially establish cell fates. One major question is whether gene expression in differentiating tissues represents a continuous trajectory versus a discontinuous progression through a series of discrete transcriptional states. Here we integrated our scRNAseq dataset with the Cao et al dataset to analyze primary notochord gene expression from its origin at the 64-cell stage through to the hatched larvae. We found that the temporal expression patterns of notochord-enriched transcripts clustered into 4-5 relatively discrete groups, supporting the idea that notochord differentiation involves a progression through several distinct transcriptional states. This concept could be developed further in the future by integrating gene expression profiling, TFBS enrichment, computational inference and perturbative experiments to build, test and refine temporally dynamic GRN models of this and other differentiating tissues.

The differentiating notochord undergoes several large-scale morphogenetic events including patterned rounds of asymmetric division, mediolateral intercalation, notochord cell elongation, notochord sheath formation and notochord lumen formation. For some of these processes, previous studies have provided insight about specific genes involved. One hypothesis is that distinct transcriptional waves in the notochord might control specific aspects of morphogenetic cell behavior. This could potentially be addressed by using Gene Ontology analysis and other strategies to assess predicted effector gene functions in the context of these different waves. These types of analyses combined with functional genomic perturbations and single-cell resolution analysis of cellular behaviors are feasible in an organ comprised of only 40 cells and would lay the groundwork for understanding how GRNs might direct morphogenesis in more complex systems.

## Author Contributions

KW: conceptualization, investigation, formal analysis, data curation, writing- original draft, writing-review and editing.

WR: investigation, writing-review and editing, supervision, project administration. MV: conceptualization, formal analysis, writing-original draft, writing-review and editing, supervision, project administration, funding acquisition.

## Acknowledgements

The authors thank the Kansas State University College of Veterinary Medicine Confocal Core, the Kansas State University Integrated Genomics Facility and the Kansas State University Beocat supercomputing cluster. We acknowledge support from the US National Science Foundation (IOS 1456555) and the US National Institutes of Health (1R01HD085909) to MV and from the Kansas State University Johnson Basic Cancer Research Center to KW.

## Materials & Methods

### Ciona

Wild caught *Ciona robusta* (formerly *Ciona intestinalis* type A) were purchased from Marine Research and Educational Products (MREP, San Diego). Animals were housed in a recirculating artificial seawater (ASW) aquarium at 18°C under constant light until experimental use.

### Experimental setup

An independent fertilization, dechorionation, dissociation, encapsulation, and library prep were performed to obtained data for each of the 64-cell, 110-cell, and mid-gastrula stages. The U0126 drug treatments were performed in parallel for each stage using embryos from the same fertilization and dechorionation.

### Fertilization, dechorionation and embryo culture

Fertilization and dechorionation were performed according to standard protocols [93]. Eggs and sperm were collected from 3-6 adults for each fertilization. Washed eggs were inseminated and left to fertilize for 5 minutes. They were then placed into dechorionation solution and gently pipetted to remove the follicle cells and chorion. Once dechorionation was complete, the eggs were washed 4 times in ASW containing 0.1% BSA. Embryos were incubated at 18°C and development was tracked in real time using in-incubator microscopy. Embryos were staged according to [45].

### Drug treatment

Embryos were treated with the MEK inhibitor U0126 (Sigma) from the 16-cell stage onward at a concentration of 4μM. At the desired stage, a 1000X stock of 4mM U0126 in DMSO was added to the culture dishes and mixed to give a final concentration of 4μM. Control embryos were treated with 0.1% DMSO at the same time as drug treated embryos.

### Dissociation

Embryos were harvested at the desired stage in a 1.5 mL microcentrifuge tube that had been treated with 1% BSA in ASW. Embryos were gently pelleted by centrifugation at 700X g for 30 seconds and the supernatant was removed. They were then washed twice with room temp Calcium-Magnesium Free AWS (CMF-ASW). Dissociation was performed by gentile pipetting of embryos in cold 0.1% BSA/CMF-ASW with 1% trypsin for 2 minutes. Following dissociation, the embryos were washed twice using cold 0.1% BSA/CMF-ASW, filtered through a 40 μm strainer and washed once more. Cells were then resuspended in cold 0.1% BSA/CMF-ASW, an aliquot was counted on a Neubauer-Improved hemocytometer, and the remainder were diluted to a concentration of 250,000 cells/mL. A small subsample of embryos from each collection was fixed in 2% paraformaldehyde in ASW to confirm staging and embryo quality by confocal microscopy.

### Encapsulation and RNA capture

Microfluidic encapsulation of single cells with barcoded beads was performed on the Dolomite Nadia instrument (Dolomite Bio, Royston UK) with a 2-lane encapsulation chip. One lane on each chip was used for the DMSO control and one for the U0126-treated sample. We used the standard Nadia protocol (v1.8) and their filter-based emulsion breaking protocol. The capture beads and chemistry used in this protocol are essentially the same as in [3] and use a stringent lysis buffer and reverse transcription after emulsion breaking. The beads are coated with oligos that incorporate a poly(T) sequence to capture mRNA 3’ ends and also two barcode sequences. One is a cell barcode that is shared between every oligo on a given bead. The other is a short Unique Molecular Identifier (UMI) barcode that differs between oligos on each bead and is used to correct for PCR duplicates in each sequenced library.

### Library Preparation and sequencing

After RNA capture and emulsion breaking, reverse transcription, exonuclease treatment, and PCR were performed according to the Dolomite protocols. Sequencing libraries were generated using the Illumina Nextera XT kit and library quality was assessed using an Agilent Bioanalyzer. Sequencing was performed on an Illumina NextSeq 500 at the KSU Integrated Genomics Facility. For all sequencing runs, an Illumina High-output 150 cycle kit was used with read lengths of: 26 bp for the Cell barcode/UMI read, 8 bp for the i7 index read, and 116 bp for the transcript read. For each timepoint, the DMSO and U0126 libraries from were sequenced in a pool across all four lanes of an Illumina chip.

### Sequence alignment and demultiplexing

Following sequencing, the dropSeqPipe pipeline [3] was used to align reads to the genome, count individual UMIs and assign them to STAMPs.

### SNP analysis

The GATK best-practices pipeline for variant calling in single-cell RNAseq data was used to call SNPs across the entire genome for all STAMPs [94]. The VCFtools package was used to calculate a relatedness statistic (unadjusted Ajk statistic based on [50]) between all pairwise STAMP comparisons in controls and drug treated embryos at each stage [95]. The resulting distance matrix was clustered using the cluster module of the python Scipy package [96].

### Dimensional reduction, clustering, and cluster analysis

Dimensionality reduction, clustering and further downstream analysis were performed using Seurat.v3 [51]. STAMPs with low numbers of genes detected (64-cell: <1750, 110-cell: <2500, mid-gastrula: <1500) were filtered out to remove likely empty droplets or low quality transcriptomes. STAMPs that were not contained in one of the well-defined “mother-of-origin” SNP clusters at each stage were also filtered out to remove potential doublets or low quality transcriptomes. For all timepoints, the default parameters for data normalization and variable feature selection were used. Each timepoint was then scaled and centered and the effects due to different mothers-of-origin were regressed out. Linear dimensional reduction was performed using PCA, and the number of PCs to keep for downstream clustering and non-linear dimensional reduction was determined by the modified Jackstraw Procedure. The number of PCs used for all downstream analysis (Non-linear dimensional reduction/UMAP, clustering, pairwise cluster distance measurement) are as follows: DMSO 64-cell – 25, U0126 64-cell – 20, DMSO & U0126 64-cell – 30, DMSO 110-cell – 30, U0126 110-cell – 25, DMSO & U0126 110-cell – 35, DMSO mid-gastrula – 35, U0126 mid-gastrula – 25, DMSO & U0126 mid-gastrula – 40, DMSO & U0126 all stages – 50. Distances between clusters were determined using the Euclidian distance between cluster centroids in a PCA reduced gene expression space that contained DMSO and U0126 treated cells at all stages.

### Cell type and lineage assignment

Cell types were assigned to clusters at each stage by identifying marker genes and validating their expression domains using the ANISEED database [46]. Lineages were assigned using the known and stereotyped lineages in the *Ciona.* Occasionally, a single cluster at one stage gave rise to more than two clusters at the next stage. When this occurred, lineages were assigned according to the known lineage relationships and these “quadfurcations” were excluded from analysis of quasi-mother/daughter relationships.

### TFBS analysis

TFBS analysis was performed similar to [74]. Control and target sequence sites were determined using the FindMarkers function implemented in Seurat to find statically significantly up and downregulated genes as desired. To extract putative enhancer regions from these gene models, we extracted the 1,500 bp of sequence upstream of the transcription start site. Additionally, we intersected these upstream regions with whole-embryo ATACseq peaks from previous experiments and kept the entire sequence of any peak that was overlapping the 1,500 bp upstream for a gene model. These sets of sequences were then saved as fasta files and used as the input for TFBS site calling and enrichment testing using the oPOSSUM3 [58] command line-tools with default settings and the JASPAR2020 vertebrate core PWMs [97].

### Differential Expression Testing

Differential expression testing was performed with the FindMarkers function implemented in Seurat using the Wilcoxon Rank-Sum test statistic. For differential expression between “sibling” clusters, testing was performed between only the two clusters. For identification of “enriched” genes in the notochord, testing was performed using the notochord cells versus all other cells.

### Average expression measurement

Pseudo-bulk expression measurements for each cluster were made using the AverageExpression function in Seurat. Pseudo-bulk counts were made for all clusters in a common gene expression space after normalization and scaling.

### Elk1/3/4 ATACseq analysis

Called peaks from Madgwick et al’s [57] early whole embryo ATACseq experiments were downloaded from ANISEED and putative TFBSs were identified using the FIMO TFBS scanner with a p-value threshold of 0.001 and the JASPAR2020 vertebrate core PWMs. Bedtools was used to associate each peak with the closest gene model in the KH2012 gene build. Sites matching the human ETS1, ELK1, ERG and ETV matrices were de-duplicated by taking the best match within a 3bp window of the center of each hit.

### Data and code availability

Raw sequencing reads, along with unprocessed (dropSeqPipe output) and processed Seurat objects for each stage are available online at GEO (accession number: in process). Code used for SNP calling and clustering as well as TFBS analysis are available on the Veeman lab github repo (https://github.com/chordmorph).

### Supplemental files and tables

Supplemental File 1: Poster image showing differential transcription factor expression and TFBS enrichment for all bifurcations.

Supplemental Table 1: Sequencing statistics

Supplemental Table 2: Number of stamps in each cluster for all treatments and developmental stages.

Supplemental Table 3: Marker genes for all clusters at the 110-cell stage

Supplemental Table 4: TFBS enrichment scores in the Neural (A) C3/4 cells

Supplemental Table 5: Transcription factor differential expression in “trios”

Supplemental Table 6: Transcription factor association with open chromatin length in *Ciona.*

Supplemental Table 7: Effector TFs associated with putative *Ciona* super-enhancers.

---

**Supplemental Figure 1: Cell lineages and division patterns are stereotyped during early *Ciona* development.** This figure summarizes the stereotyped cell lineages of early ascidian embryos based on the work of ([18–21,41,98,99]). A-D’) Animal and vegetal views of the *Ciona* embryo at the 16-cell (A-A’), 32-cell (B-B’), 64-cell (C-C’), and 110-cell (D-D’) stages. On each view, the blastomere name is labeled on the right half of the embryo, and the cell type is colored on the left half of the embryo for lineage restricted blastomeres. E-E’) Fate map of the animal blastomeres at the 110-cell stage (E), and their future territory at the mid-tailbud stages (E’). Colors represent sibling cell pairs. Adapted from [19]. F) Cartoon representation of the neural plate at mid-gastrula stage. Blastomere names are printed on the right half of the neural plate. Arabic numerals define the column numbers, while roman numerals define the row numbers. Colors on the left half of the neural plate define the cellular lineage and correspond with those in A-D’ and G (magenta: A-line, orange: a-line, brown: b-line). G) Lineage tree of blastomeres in the early stages of *Ciona* development. Colors correspond with A-D’ and F.

**Supplemental Figure 2: Adult of origin effect correction using SNPs.** A-C) First-pass UMAP plots at the 64-cell (A), 110-cell (B), and mid-gastrula (C) stages where the animal lineages are colored in red. D-F) Dendrograms of heterarchical clustering of genetic relatedness of STAMPs in the 64-cell (D), 110-cell (E), and mid-gastrula (F) stages using SNPs. G-I) UMAP plots at the 64-cell (G), 110-cell (H), and mid-gastrula (I) stages where STAMPs have been colored by the putative adult-of-origin using the clustering in D-F. UMAP space is identical to A-C.

**Supplemental Figure 3: The secondary notochord also exhibits transcriptional waves.** A-B) Temporal expression profiles of all secondary notochord enriched genes (A), and transcription factors (B) reveal distinct waves of expression throughout the course of development similar to the primary notochord.

**Supplemental Figure 4: TFBS enrichment in the primary notochord**. Left panel) Average TFBS motif enrichment z-score in the primary and secondary notochord from the 64-cell, 110-cell, and mid-gastrula stages, which represent the early notochord GRN. Right panel) Average TFBS motif enrichment z-score in the primary and secondary notochord from the final three stages of development assayed, which represent the late notochord GRN.

