## Supplementary figures and images for "Single-cell analysis of cell fate bifurcation in the chordate *Ciona*"

### Supplemental File 1

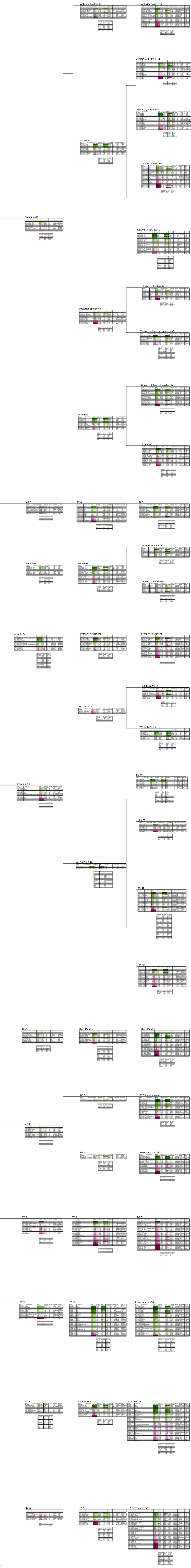
