## Supplemental Figures for "Single-cell analysis of cell fate bifurcation in the chordate *Ciona*"

### Supplemental Figure 1

#### Cell lineages and division patterns are stereotyped during early Ciona development.

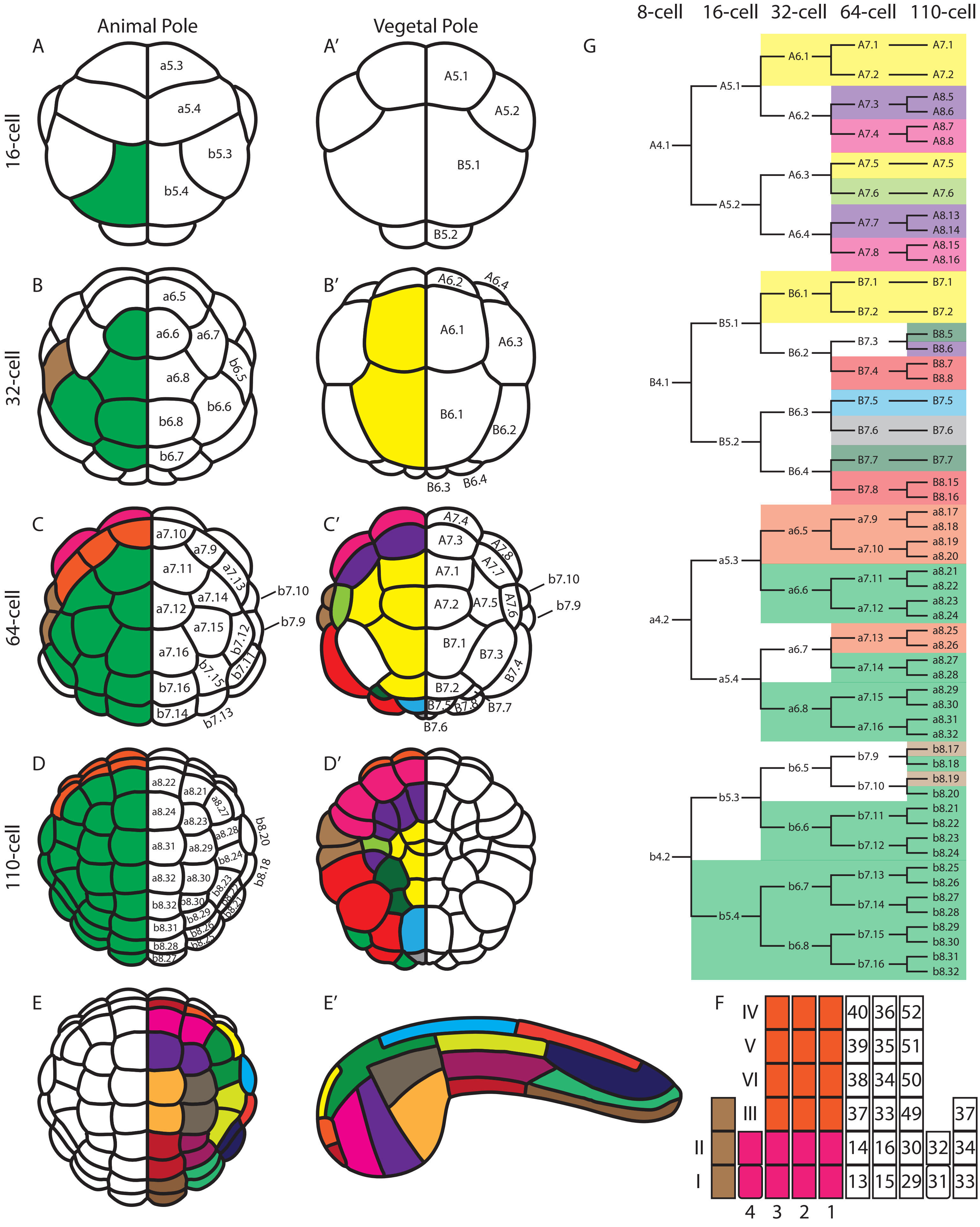

### Supplemental Figure 2

#### Adult of origin effect correction using SNPs.

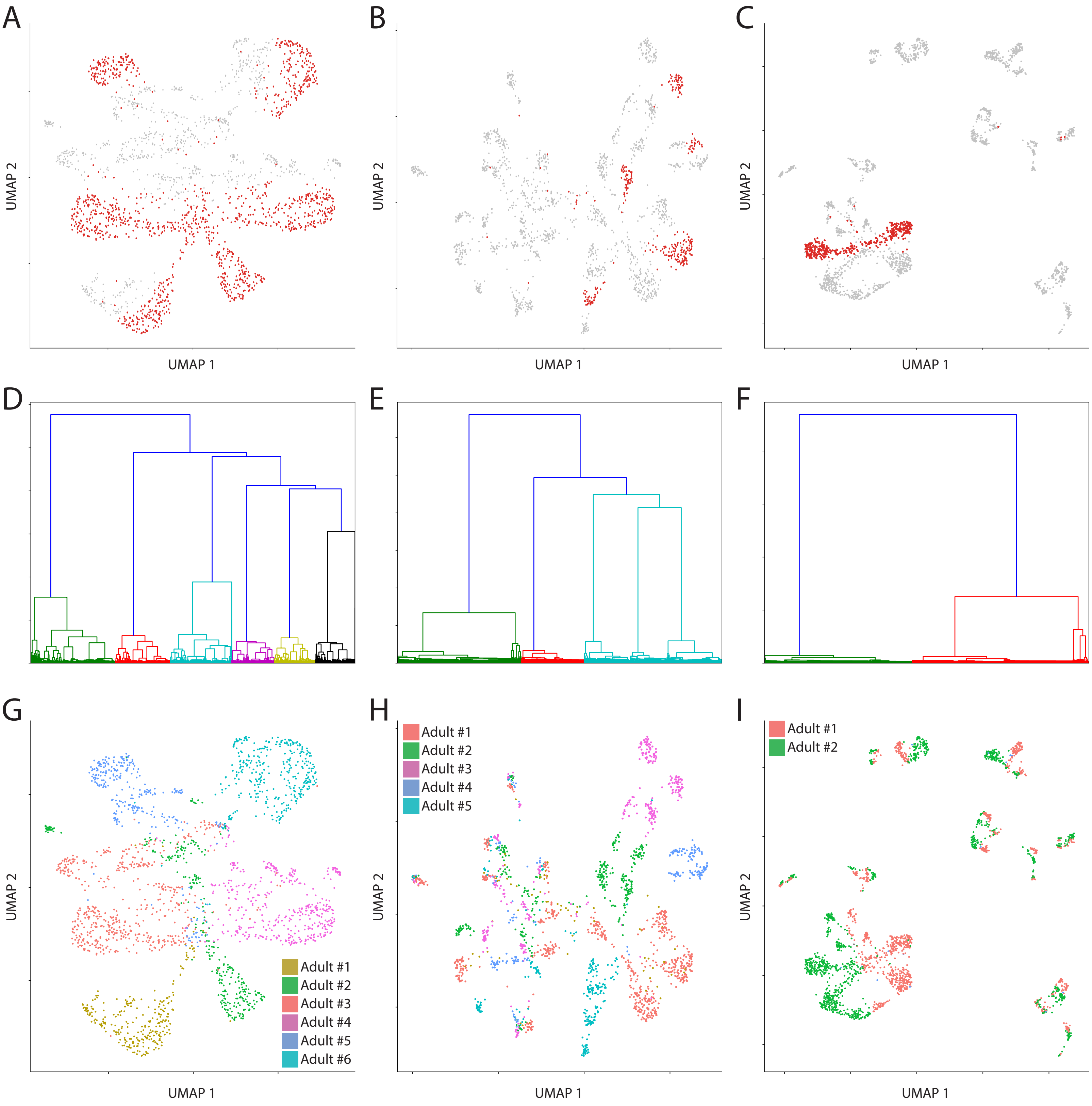

Supplemental Figure 3

The secondary notochord also exhibits transcriptional waves.

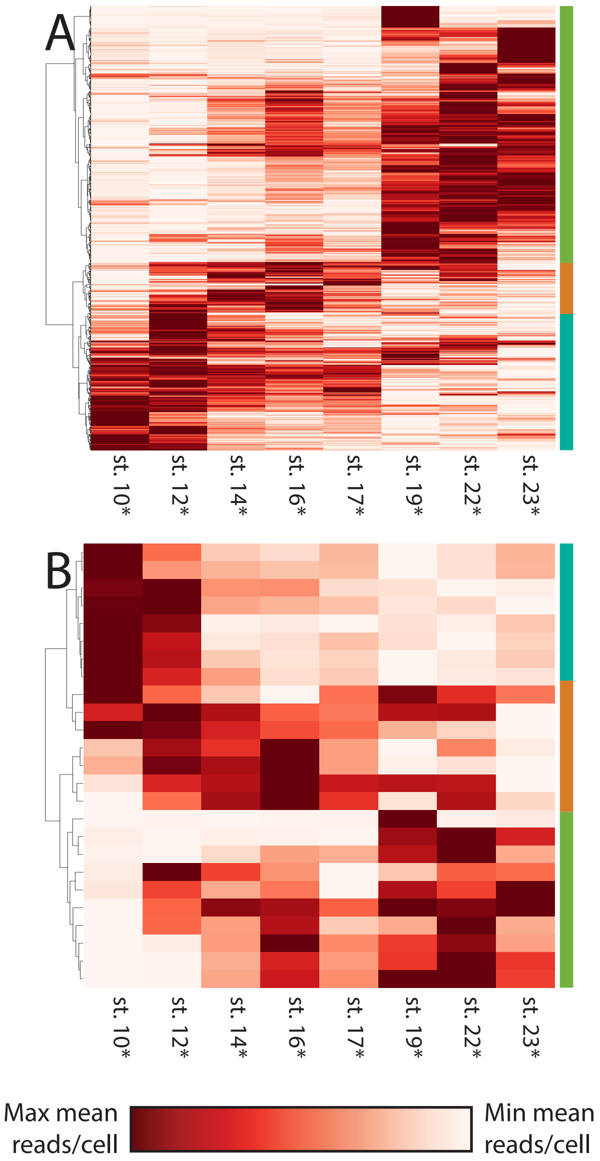

### Supplemental Figure 4
TFBS enrichment in the primary notochord.

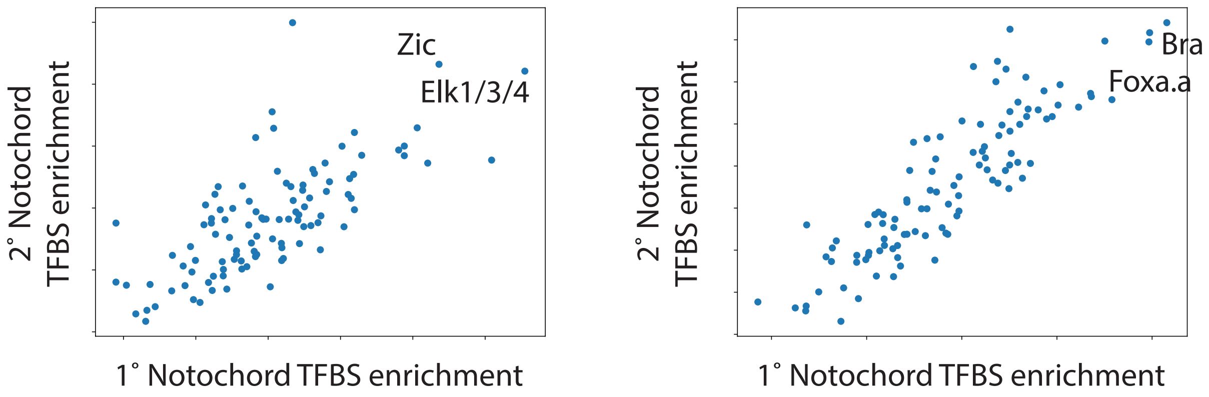
